# Characterizing heterogeneity along EMT and metabolic axes in colorectal cancer reveals underlying consensus molecular subtype-specific trends

**DOI:** 10.1101/2023.08.06.551165

**Authors:** Manas Sehgal, Soundharya R, Joel Markus Vaz, Raja G Yogeshwar, Srinath Muralidharan, Sankalpa Venkatraghavan, Mohit Kumar Jolly

## Abstract

Colorectal cancer (CRC) is highly heterogenous with variable survival outcomes and therapeutic vulnerabilities. A commonly used classification system in CRC is the Consensus Molecular Subtypes (CMS) based on gene expression patterns. However, how these CMS categories connect to axes of phenotypic plasticity and heterogeneity remains unclear. Here, we analyze 101 bulk transcriptomic datasets, along with patient tumor samples from TCGA and single-cell RNA sequencing data, to evaluate the extent of variation among CMS subtypes across metabolic plasticity and EMT axes. Our results show that CMS2 and CMS3 samples were relatively more epithelial as compared to CMS1 and CMS4. Single-cell RNA-seq analysis of CMS1 revealed two subpopulations: one close to CMS4 (more mesenchymal) and the other closer to CMS2 or CMS3 (more epithelial), indicating a partial EMT-like behavior. Further, in our meta-analysis and in TCGA data, epithelial phenotype score was positively correlated with scores of glycolysis, OXPHOS and FAO pathways, while mesenchymal scores showed CMS subtype-specific associations with metabolic axes. PD-L1 activity scores, however, consistently correlated positively with mesenchymal signature ones and negatively with epithelial signature ones, across the four CMS categories. Together, our results quantify the patterns of two interconnected axes of phenotypic heterogeneity - EMT and metabolic reprogramming - at a CMS subtype level in CRC.

## Introduction

Colorectal cancer (CRC) is a multifaceted malignancy that arises from the epithelial lining of the colon or rectum (Ferlay et al. 2019). It is the third most prevalent cancer worldwide (de Abreu et al. 2023). Although surgical resection is the primary method of treatment for localized tumors, non-resectable tumors present substantial clinical challenges (Sun et al. 2019). 5-fluorouracil (5-FU) was the first effective chemotherapy for CRC. Targeted treatments such as cetuximab and panitumumab (that block EGFR), bevacizumab (that prevents angiogenesis by blocking VEGF), pembrolizumab (that blocks the immune checkpoint molecule PD-1) and vemurafenib (that inactivates BRAF V600 kinase) are frequently used (Piawah and Venook 2019; El Bali et al. 2021). The inherent diversity in molecular and phenotypic heterogeneity of tumors leads to variable responses to targeted therapies and survival outcomes (Xie et al. 2020; Chowdhury et al. 2021; Deshmukh et al. 2023).

To characterize this diversity, the Consensus Molecular Subtypes (CMS) system, a classification of CRC tumors based on the transcriptomic profile (Guinney et al. 2015) was proposed. Based on gene expression profiles, the ‘CMSclassifier’ divided the CRC tumors into four subtypes, using a random forest algorithm based on gene expression signals from tumor’s immune and stromal compartments: a) Microsatellite instability-immune, or CMS1 tumors, that show significant immune activation and genomic instability, b) ‘Canonical’ CMS2 tumors that exhibit WNT and MYC signalling pathway activation, c) ‘Metabolic’ CMS3 tumors that show Epithelial-mesenchymal transition (EMT) features and metabolic dysregulation, and d) ‘Mesenchymal’ CMS4 tumours that display significant stromal infiltration, angiogenesis, and mesenchymal characteristics (Fessler and Medema 2016; Menter et al. 2019; Eide et al. 2021). However, CMSclassifier algorithm sometimes failed to correctly identify CMS4-mesenchymal sub-population in cell lines, patient-derived organoids, and xenografts. Thus, CMSCaller, a more robust classifier developed lately, is being increasingly adopted to deliver a more accurate subtype classification utilizing multiple sources of transcriptomic data (Eide et al. 2017).

CMS classification provides valuable therapeutic insights. For instance, clinical studies have found that the CMS1 patients who received bevacizumab had significantly higher overall survival and progression free survival rates compared to CMS1 patients who received cetuximab (Stintzing et al. 2019; Rebersek 2020) whereas for CMS4 tumors, irinotecan (IRI)-based chemotherapy outperforms oxaliplatin (OX)-based chemotherapy (Okita et al. 2018). Since each subtype responds differently to therapies, CMS classification offers more personalized strategies for therapeutic interventions, thus improving treatment outcomes.

Recently, another method is being increasingly used to quantify the extent of heterogeneity in tumor cells using single-cell transcriptomic data (scRNA-seq) - Shannon Entropy. Entropy is derived from information theory and is helpful in providing a quantitative measure of the diversity and distribution of gene expression within cell populations (Gandrillon et al. 2021; García-Nieto et al. 2022). Moreover, Shannon entropy can be used to identify key genes driving cellular heterogeneity. Genes with high entropy are often associated with the dynamics of lineage specification and cell-fate decisions. Their expression patterns indicate transcriptional plasticity and cell-state transitions, providing valuable insights into underlying cellular dynamics driven by regulatory networks (Kharchenko, 2021; Berretta & Moscato, 2010). Thus, investigating the entropy of tumor cells, along with their functional attributes, is important to fully understand the underlying biological variability and vulnerabilities in CRC.

Here, we analyze 101 bulk transcriptomic datasets, along with patient tumor samples in colorectal cancer from The Cancer Genome Atlas (TCGA) and single-cell RNA sequencing data, to evaluate the degree of variation among CMS subtypes across metabolic reprogramming and EMT axes. Further, we analyzed the consequences of associations between these axes for disease prognosis. Our results show that the epithelial phenotype score was positively correlated with scores of glycolysis, OXPHOS and FAO pathways, while mesenchymal scores showed CMS subtype-specific associations with metabolic axes. PD-L1 activity scores consistently correlated positively with mesenchymal signature ones and negatively with epithelial signature ones, across CMS categories. Finally, we observed that CMS2 and CMS3 were more epithelial as compared to CMS1 and CMS4. Interestingly, single-cell RNA-seq data revealed two subpopulations in CMS1: one close to CMS4 (more mesenchymal) and the other closer to CMS2 or CMS3 (more epithelial), indicating a partial EMT-like behavior. Together, our results quantify the patterns of epithelial-mesenchymal heterogeneity and interplay between EMT and metabolic plasticity at CMS subtype level in CRC.

## Methods

### Software and Datasets

Computational and statistical analyses were conducted using R (version 4.3.0) and Python (version 3.9). Microarray datasets were retrieved from National Center for Biotechnology Information’s Gene expression omnibus (NCBI GEO) using the ‘GEOquery’ R package. Processed RNA sequencing and single-cell RNA sequencing data were also obtained directly from individual datasets from the NCBI GEO database (**Table S1**). TCGA datasets were obtained using UCSC Xena tools for COAD_READ.

### Pre-processing of datasets

Pre-processing of microarray datasets was conducted to obtain the gene-wise expression from the probe-wise expression matrix using respective annotation files for the mapping of probes to genes. In case multiple probes were mapped to a single gene, the mean expression of all mapped probes was utilized to obtain the final values for those genes.

Raw counts obtained for RNA and single-cell RNA sequencing data were normalized for gene length and transformed to transcripts per million (TPM) values. They were then log2 normalized to acquire the final expression data.

For single-cell RNA sequencing (scRNA-seq) datasets, MAGIC (version 2.0.3) (van Dijk et al. 2018). imputation algorithm was utilized to recover noisy and sparse single-cell data using diffusion geometry. To map individual reads to corresponding genes, relevant platform annotation files were utilized.

### CMS Classification

CMS classification for colorectal cancer samples and tumor cells was carried out using ‘CMScaller’ (Eide et al. 2017). ‘CMScaller’ uses the nearest template prediction algorithm to assign the CMS (CMS1-CMS4) to each sample with a prediction distance with a corresponding p-value. The predictions with insignificant p-values (p > 0.05) are not assigned any subtype. The CMS template genes were also used as signatures for ssGSEA scoring to identify the enrichment of the CMS-specific signatures in the sample.

### Gene signature scoring

To quantify the enrichment of epithelial and mesenchymal signatures independently, ssGSEA (single sample gene set enrichment analysis) was performed on epithelial (for Epi scores) and mesenchymal (for Mes scores) gene signatures (Tan et al. 2014) separately using GSEAPY python library for bulk RNA sequencing and microarray datasets. Normalized enrichment score (NES) for these genesets was obtained for further analysis. A higher NES score corresponds to enrichment of that particular phenotype for the given sample. Similarly, the gene signatures for hallmark EMT, hallmark Fatty Acid Oxidation, and hallmark Glycolysis were obtained from molecular signatures database MSigDB and the respective scores were calculated (Liberzon et al. 2011). PD-L1 signature was curated as reported earlier (Sahoo et al. 2021), wherein the top correlated genes (Spearman correlation coefficient > 0.5 and p < 0.01) with PD-L1 levels in at least any 13 out of 27 cancer types were considered (**Table S2**).

The activity scores for metabolic pathways, PD-L1, and E/M signatures for single-cell RNA sequencing datasets were computed using AUCell (version 1.18.1) (Aibar et al. 2017) from the ‘Bioconductor’ package (Gentleman et al. 2004) in R package with default parameters.

### Survival analysis

Survival data were obtained from the TCGA cohort of patients for colorectal cancer. The samples were categorized into CMS high, and CMS low groups based on median of the respective CMS scores for each CMS subtype. Kaplan-Meier analysis was performed using the R package ‘survival.’ A log-rank test was used to compute the *p*-values. The reported hazard ratio (HR) and confidence interval (95% CI) were determined using Cox regression using the ‘coxph’ function.

Additionally, colorectal cancer samples were also split into High (+) and Low (-) expression subgroups based on the median for different gene signatures (Epi, Mes, FAO, glycolysis, OXPHOS, and PD-L1) scores. The effect of the simultaneous enrichment on survival was calculated for all gene signatures in a pairwise manner. HR and p-values were depicted in forest plots created using the ‘ggforest’ function from the ‘survminer’ package.

### Entropy Calculation

Cell-wise entropy values were calculated for each geneset using the following formula (Kannan et al. 2021):

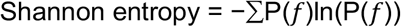

Where P(*f*) is the ratio of the normalized expression value of a particular gene to the sum of all gene expression values for a cell. The entropy values were normalized for the number of genes in each gene set to maintain comparability.

## Code availability

Codes used in this study are available at https://github.com/Soundharya-R/CRC

## Data availability

Publicly available transcriptomics datasets from NCBI GEO (**Table S1**), TCGA cohorts from UCSC Xena and single-cell RNA sequencing data (GSE132465, GSE144375) were analyzed in this study.

## Results

### 3.1 Associations between EMT, metabolism axes and immune evasion and their associations with patient survival

To assess the associations between multiple axes of plasticity governing CRC heterogeneity, we began by investigating three important aspects - cancer cell metabolism, immune evasion, and EMT. We investigated the associations among these three axes in 101 bulk transcriptomic datasets with CRC samples (**Table S1**) by calculating ssGSEA scores of associated gene signatures. We observed that among the 43 cases where the epithelial phenotype correlated significantly (p < 0.05) with the PD-L1 pathway, the correlation coefficient was negative in 36 cases, indicating that epithelial phenotype is largely negatively correlated with immune evasion (**Fig 1A**). Conversely, mesenchymal phenotype predominantly correlated positively with the PD-L1 signature (60 out of 61 cases). Epithelial and mesenchymal signatures showed a dominantly negative correlation with each other (53 out of 58 datasets), as expected (**Fig S1**). Similar antagonistic trends were seen for epithelial vs. mesenchymal signatures with the metabolic axes - while the epithelial phenotype correlated positively with oxidative phosphorylation (OXPHOS) (47 out 51 datasets), and fatty acid oxidation (FAO) (65 out of 67 datasets), mesenchymal phenotype enrichment was negatively correlated with OXPHOS (42 out of 48 datasets) and FAO axes (25 out of 31 datasets) (**Fig 1B**). The trend with the glycolysis signatures was not as strong, yet antagonistic for the epithelial vs. mesenchymal signature ssGSEA scores. These trends are reminiscent of our pan-cancer analysis showing that FAO and OXPHOS were negatively correlated with EMT (Muralidharan et al., 2022). However, we had noticed an association between glycolysis with partial or full EMT, which is not as strongly recapitulated in CRC data. Such context-specific differences may emerge from the heterogeneity in CRC samples from a CMS context.

**Figure 1:**
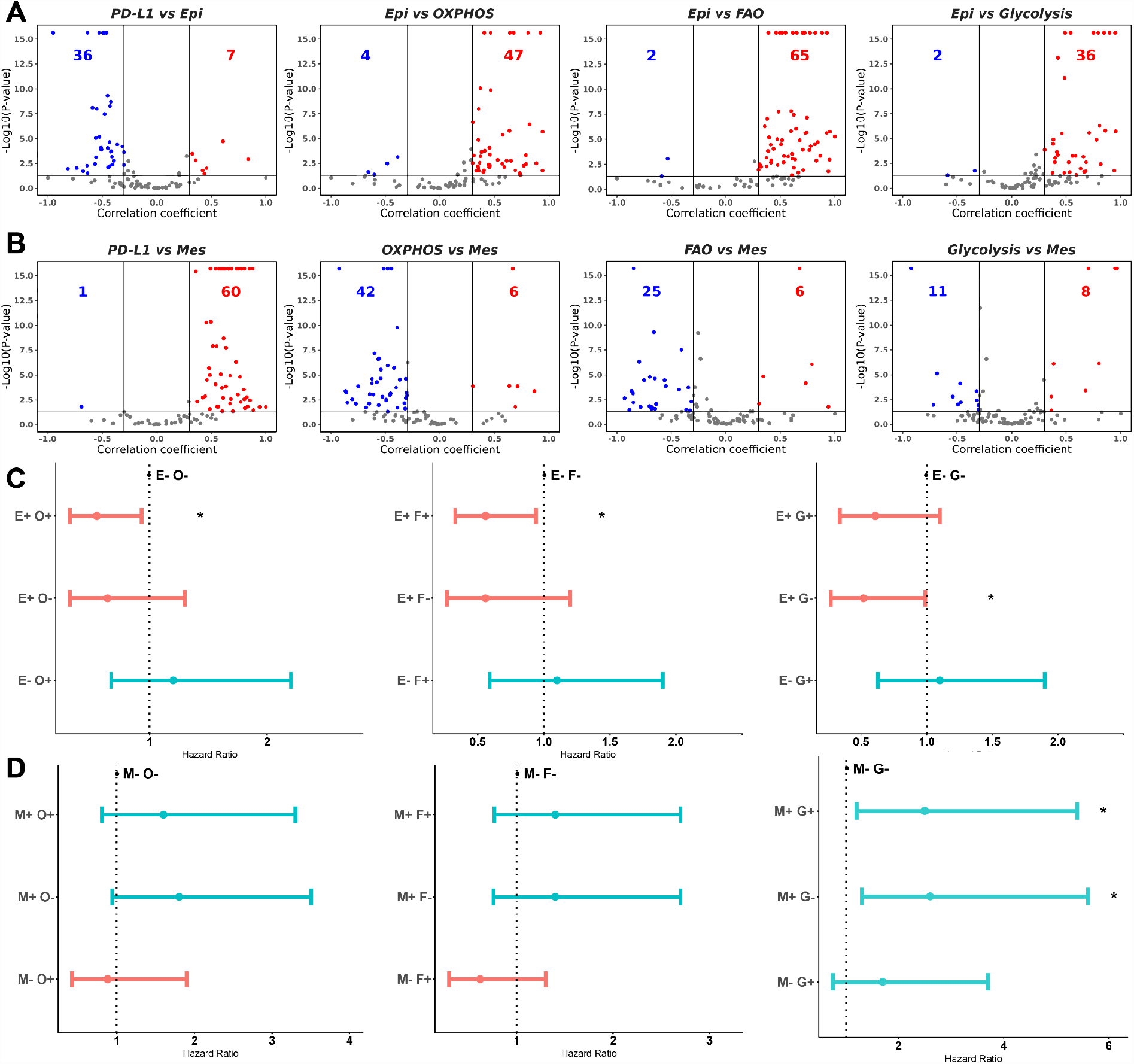
Relationship between EMT, metabolism, and PD-L1 signatures in CRC bulk-level transcriptomes and differences in survival probabilities for their pairwise concurrent enrichment. **A)** Volcano plots depicting Spearman correlation coefficient (x-axis) and -log10(p-values) (y-axis) for PD-L1 vs. Epi scores (left), Epi vs. OXPHOS (middle-left), Epi vs. FAO (middle-right) and Epi vs Glycolysis scores (right). Boundaries for significant correlation are set at R > ± 0.3 and p < 0.05. Red data points indicate datasets for which the association is significantly positive, blue for negative, and gray for insignificant correlation. **B)** Same as A) but for PD-L1 vs. Mes (left), OXPHOS vs. Mes (middle-left), FAO vs. Mes (middle-right), and Glycolysis vs. Mes scores (right). **C)** Forest plots depicting mean hazard ratios (HR) ± 95% confidence intervals and corresponding p-values (‘*’ for p <0.05) for overall survival associated with simultaneous enrichment of epithelial and OXPHOS (left), epithelial and FAO (middle) and epithelial and Glycolysis signatures (right). (+) and (-) subgroups are based on median values. Mean HR values > 1 are shown in blue, while those < 1 are shown in red. same as C) but for mesenchymal and OXPHOS (left), mesenchymal and FAO (middle), and mesenchymal and Glycolysis signatures (right).

Next, to test the clinical outcomes of simultaneous enrichment of these metabolic pathways and PD-L1 with epithelial and mesenchymal phenotypes, we performed survival analysis on the colorectal cancer patient data in TCGA database. Our evaluation of these pairwise comparisons revealed that upregulation of epithelial signature along with higher expression of FAO and OXPHOS genes (E+F+ and E+O+) had significantly better overall survival probability compared to Epi-Low/FAO low group (E-F-) and Epi-Low/OXPHOS-low group (E-O-) respectively. No such trend was observed for glycolysis signature, though (**Fig 1C**). In contrast, the combination of high expression of glycolysis pathway genes with the mesenchymal phenotype (G+M+) has significantly worse implications for survival probability with reference (G-M-) (**Fig1D**), but no trend was noticed for co-enrichment of mesenchymal with FAO or OXPHOS. Additionally, the PD-L1 signature, despite being positively correlated with mesenchymal signatures, is associated with better survival probability in CRC (**Fig S1**). This analysis emphasizes the strong association of epithelial phenotype with a better survival response, whereas the presence of mesenchymal phenotype is generally associated with a worse prognosis. Further, the metabolic pathways, such as FAO and OXPHOS, that correlated positively with epithelial phenotype, had a similar effect on the hazard ratio (HR). On the contrary, the glycolysis signature and PD-L1 signature, although more strongly positively associated with epithelial phenotype and mesenchymal phenotype respectively, tend to affect the hazard ratios in opposite directions. Together, these results suggest an interplay between the different interconnected axes of cellular plasticity (EMT, metabolic switching) in mediating patient survival (Jia et al. 2021).

### 3.2 Heterogeneity among Consensus Molecular Subtypes (CMS) and their associations with EMT and different axes of metabolism and immune evasion

The intra-tumor heterogeneity in CRC patients can be better studied by subdividing cancer samples into subtypes based on their shared features and differences at the molecular and cellular levels. The subtypes differ widely in their transcriptomic and genomic profiles while also showing differences in prevalence in parts of the colon (Chowdhury et al. 2021).

To begin, we performed Kaplan Meier survival analysis to determine the clinical outcome for the enrichment of different CMS using patient data from TCGA database. The CMS scores were calculated with ssGSEA using the subtype-specific template signatures used by ‘CMSCaller’ function. We hypothesized that the samples with higher CMS2 and CMS3 scores would have better survival probabilities than those with higher CMS4 scores since the latter has upregulation of EMT pathway, correlating to poor patient outcomes (Matsuyama et al. 2019).

The Kaplan Meier plots compared CMS-high versus CMS-low expression for the four subtypes, and we observed the hazard ratios for samples with high CMS1 and 4 scores were lower than 1 (i.e. enrichment of CMS1 or CMS4 associated with worse survival), and hazard ratios for samples with higher CMS2 and 3 scores were lower than 1. (i.e., enrichment of CMS2 or CMS3 associated with better survival) (**Fig 2A**). However, the trend was statistically significant only for CMS2 and CMS4.

**Figure 2:**
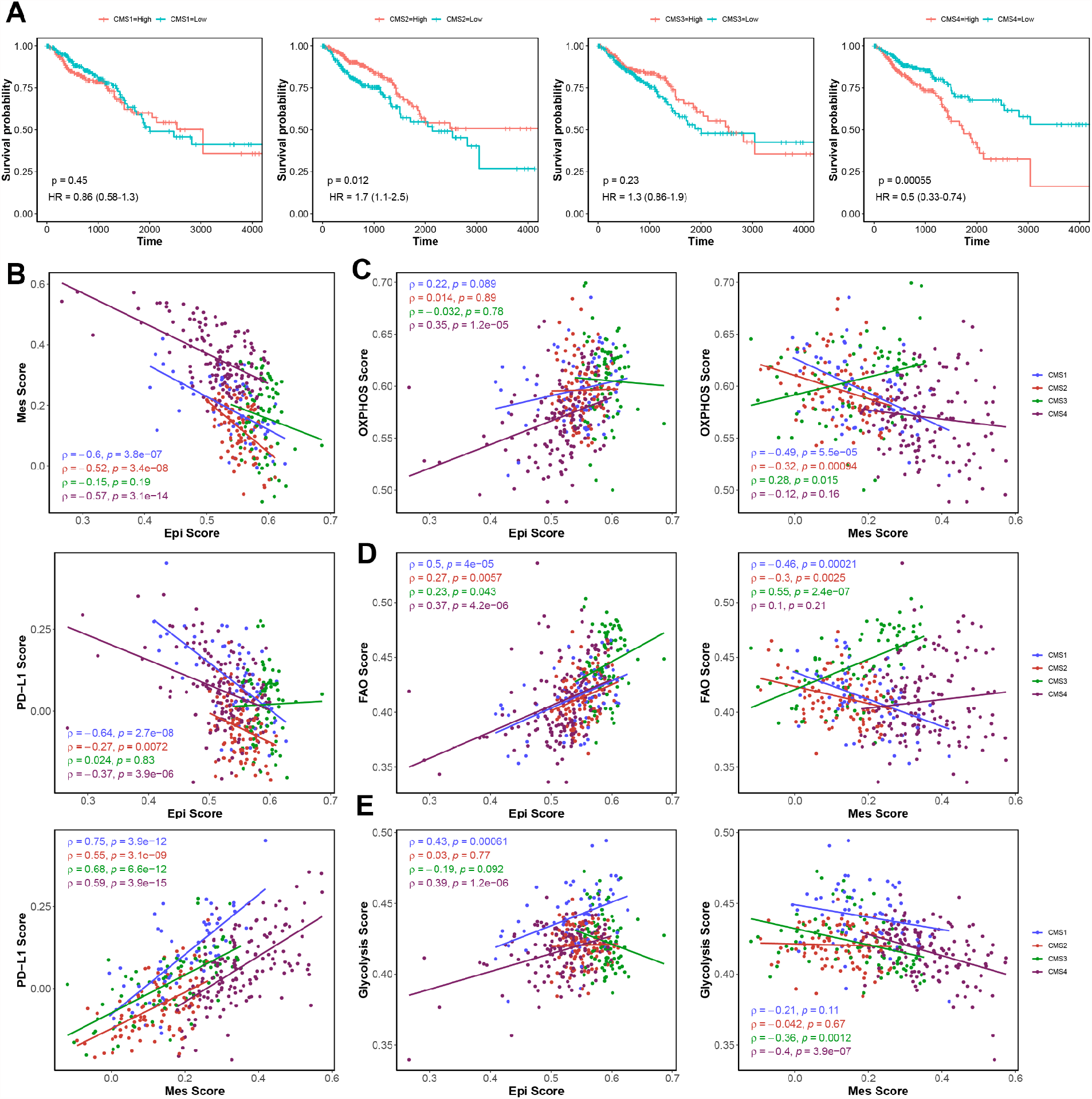
CMS-specific differences in survival probabilities and associations between EMT, metabolism, and PD-L1 in TCGA colorectal cancer samples. **A)** Kaplan-Meier curves showing differences in survival probabilities for CMS1-high (red) and CMS1-low (blue) (left), CMS2-high and low (middle-left), CMS3-high and low (middle-right) and CMS4-high and low (right). Reported p-values are based on a log-rank test and indicate differences in survival between the subgroups. Mean hazard ratios (HR) ± 95% confidence intervals (95% CI) are shown. **B)** Scatter plot illustrating subtype-wise Epi (x-axis) and Mes (y-axis) scores (top), Epi and PD-L1 scores (middle), and Mes and PD-L1 scores (bottom) for TCGA colorectal cancer patient samples. Pearson’s correlation coefficient ‘ρ’ and p-values for subtype-wise correlation are shown. Blue data points are for cells classified as CMS1, red for CMS2, green for CMS3, purple for CMS4. **C)** same as **B)** but for Epi and OXPHOS scores (left) and Mes and OXPHOS scores (right), **D)** Epi and FAO scores (left) and Mes and FAO scores (right), and **(E)** Epi and Glycolysis scores (left) and Mes and Glycolysis scores (right).

To further understand the variability among the CMS subtypes, we looked at the correlation between well-studied signatures pertaining to metabolism, immune evasion, and EMT, in TCGA data available for CRC, in a CMS-specific manner (**Fig 2**). We observed that when CRC samples are not segregated by CMS, the common trends seen for the bulk dataset meta-analysis (**Fig 1**) hold true, i.e., a positive correlation between epithelial scores with FAO, OXPHOS and glycolysis scores, a positive correlation between mesenchymal and PD-L1 scores, and a negative correlation for Epi scores vs. Mes scores, and a negative correlation of Epi scores with PD-L1 scores (**Fig S2**). However, as we delved into examining these trends at the CMS subtype level, we saw CMS subtype-specific differences. For instance, in the case of glycolysis scores versus epithelial and mesenchymal scores (**Fig 2E**), the variability in the correlation coefficient values seen (both positive and negative values across 4 CMS subtypes) seem to explain their seemingly counterintuitive association with respect to epithelial and mesenchymal scores seen in the bulk data analysis noted earlier (**Fig 1A-B)**. We further noticed that epithelial and mesenchymal programs were negatively correlated even at individual subtype-level and PD-L1 signature scores associated positively with mesenchymal ones across the CMS subtypes. While, in CMS3 samples, the PD-L1 and Epi scores did not show any association, in the rest of the subtypes they were negatively correlated (**Fig 2B**). OXPHOS was shown to be positively linked with Epi and negatively associated with Mes in CMS1, CMS2 and CMS4 but these associations were found to be in the reverse direction in the CMS3 samples (**Fig 2C**). Similarly, in CMS3 and CMS4, FAO and Mes are correlated positively while in CMS1 and CMS2, they are negatively associated. However, FAO was associated positively with epithelial phenotype in all four subtypes to a similar extent (**Fig 2D**). Such differences in the relationship between these two key axes of plasticity (EMT, metabolic switching) at subtype level could serve as a distinguishing functional role of different CMS categories.

### 3.3 Different CMS subtype samples have varied status of epithelial-mesenchymal plasticity

Given the observed role of epithelial vs. mesenchymal phenotypes in determining patient survival, we assessed the extent of epithelial and mesenchymal enrichment across CMS subtypes. We chose five datasets with samples corresponding to each subtype (GSE196576, GSE161158, GSE96528, GSE14333, GSE14095). The samples in each dataset were classified into the respective subtypes using the ‘CMSCaller’ package. Here, we consistently observed that the Epi score was the highest in CMS3 subtype, followed by CMS2, whereas the Mes scores were comparable for these two subtypes. Further, we saw that CMS4 samples had the highest Mes score and lowest Epi score among all and was followed by CMS1 samples (**Fig 3A**).

**Figure 3:**
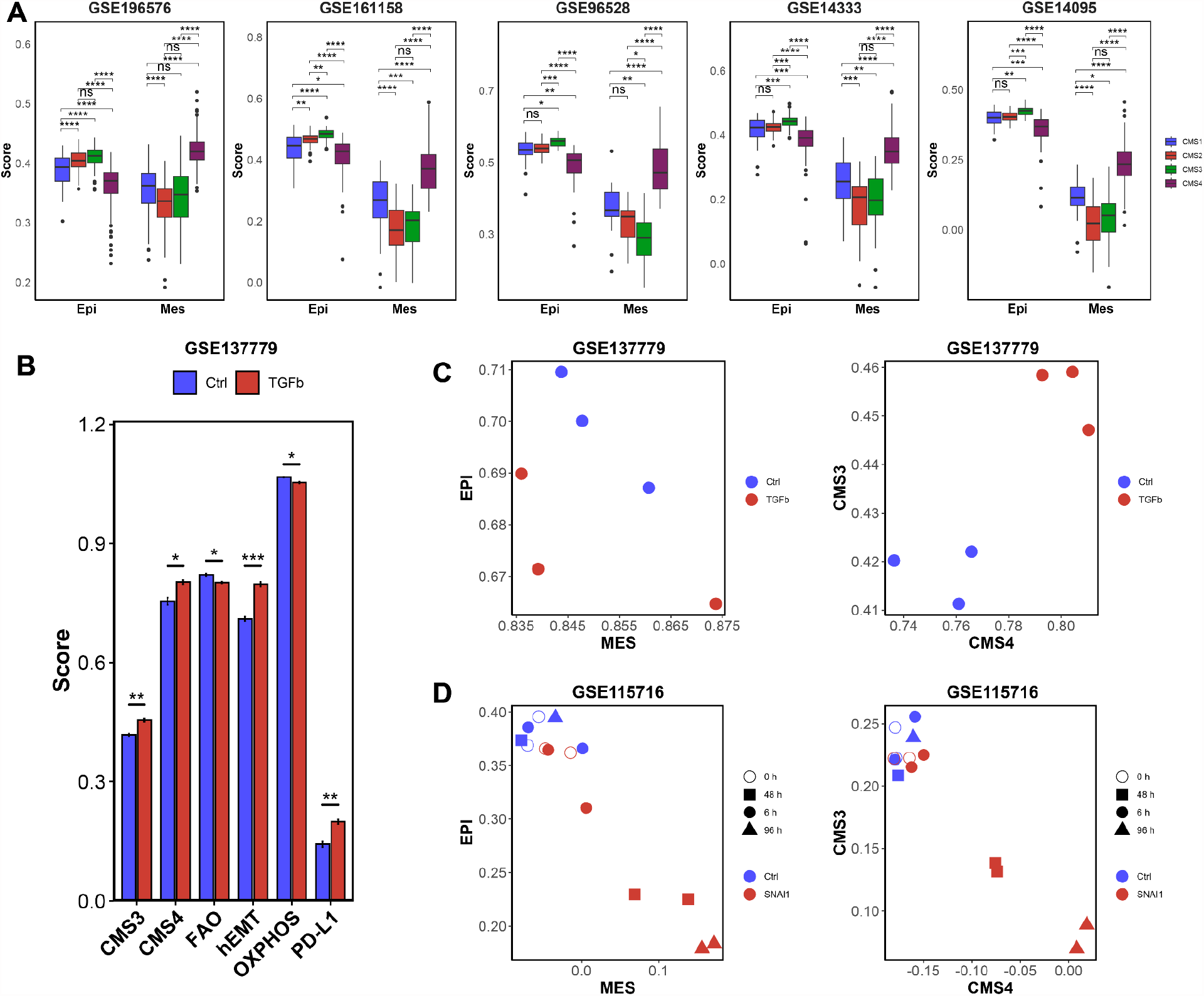
Relationship between CMS and EMT induction in bulk datasets. **A)** Boxplots showing differences in epithelial and mesenchymal scores in a subtype-specific manner. *, **, ***, **** denote p < 0.05, 0.01, 0.001, 0.0001, respectively. ‘ns’ indicates p>0.05 for student’s two-tailed t-test with unequal variance. Plot titles denote NCBI-GEO dataset IDs. **B)** Barplots showing ssGSEA scores of CMS3, CMS4 genes, FAO, hallmark EMT, OXPHOS, and PD-L1 gene sets in control (blue) vs. TGFβ-treated samples (red). **C)** Scatterplot with Epi (y-axis) and Mes (x-axis) scores (left) and CMS3 (y-axis) and CMS4 (x-axis) scores of control and TGFβ-treated samples of GSE137779. **D)** Same as **C)** but for control samples and samples with SNA1 overexpression (GSE115716).

Next, we obtained two datasets, one with TGFβ treated samples (GSE137779) and the other having samples with overexpression of EMT-inducing transcription factor SNAI1 over time (GSE115716) to gauge the extent of influence of EMT on the CMS sub-classification score. We compared the changes in CMS3 and CMS4 when samples were treated with TGFβ or SNAI1. Expectedly, we observed that SNAI1 overexpression causes a decrease in Epi scores and an increase in Mes scores. Intriguingly, we observed that SNAI1 overexpression led to higher CMS4 enrichment scores but reduced CMS3 scores (**Fig 3D**), indicating that SNAI1, a potent EMT inducer, possibly plays a role in controlling CMS plasticity. On the other hand, while TGFβ treatment reduced the Epi scores, Mes scores do not show the expected increase. While projecting the same set of samples on the CMS3/CMS4 axes, we saw that TGFβ seemed to increase both CMS3 as well as CMS4 scores (**Fig 3C**). Apart from this, we also saw that TGFβ significantly decreases FAO and OXPHOS and increases PD-L1 activity (**Fig 3B**). These examples illustrate that the mode of EMT induction dictates the extent of EMT observed as well as corresponding changes in their molecular subtyping. Another reason for this difference can be non-EMT associated changes driven by TGFβ in cellular response.

### 3.4 Single-cell RNA-sequencing analysis reveals CMS subtype-specific patterns of epithelial-mesenchymal heterogeneity

Our bulk-level CMS-specific investigation highlighted associations between CMS subtyping and EMT. We examined these associations at individual cell level through single-cell RNA-sequencing (scRNA-seq) datasets GSE132465 and GSE144375 (Lee et al. 2020). The scRNA-seq data were filtered for tumor cells, and ‘CMSCaller’ was used to assign the appropriate CMS subtype. Only the statistically significant predictions in the context of CMS assignment were used for further analysis. We noticed all the four CMS subtypes to be well-represented in these two scRNA-seq datasets (**Fig 4A, i-ii**). First, we observed that the antagonism between epithelial and mesenchymal axes is maintained even at the single-cell level (**Fig 4B, ii-ii**) across cells belonging to all four CMS subtypes. Interestingly, we also noticed a spectrum of epithelial-mesenchymal states in this two-dimensional projection: while cells classified to belong to CMS4 cells localized in (high mesenchymal, low epithelial) area, the ones classified as CMS2 and CMS3 were centered around (low mesenchymal, high epithelial) area. The cells categorized as CMS1 occupied intermediary position, indicating a hybrid E/M phenotype with the simultaneous enrichment of both epithelial and mesenchymal characteristics.

**Figure 4:**
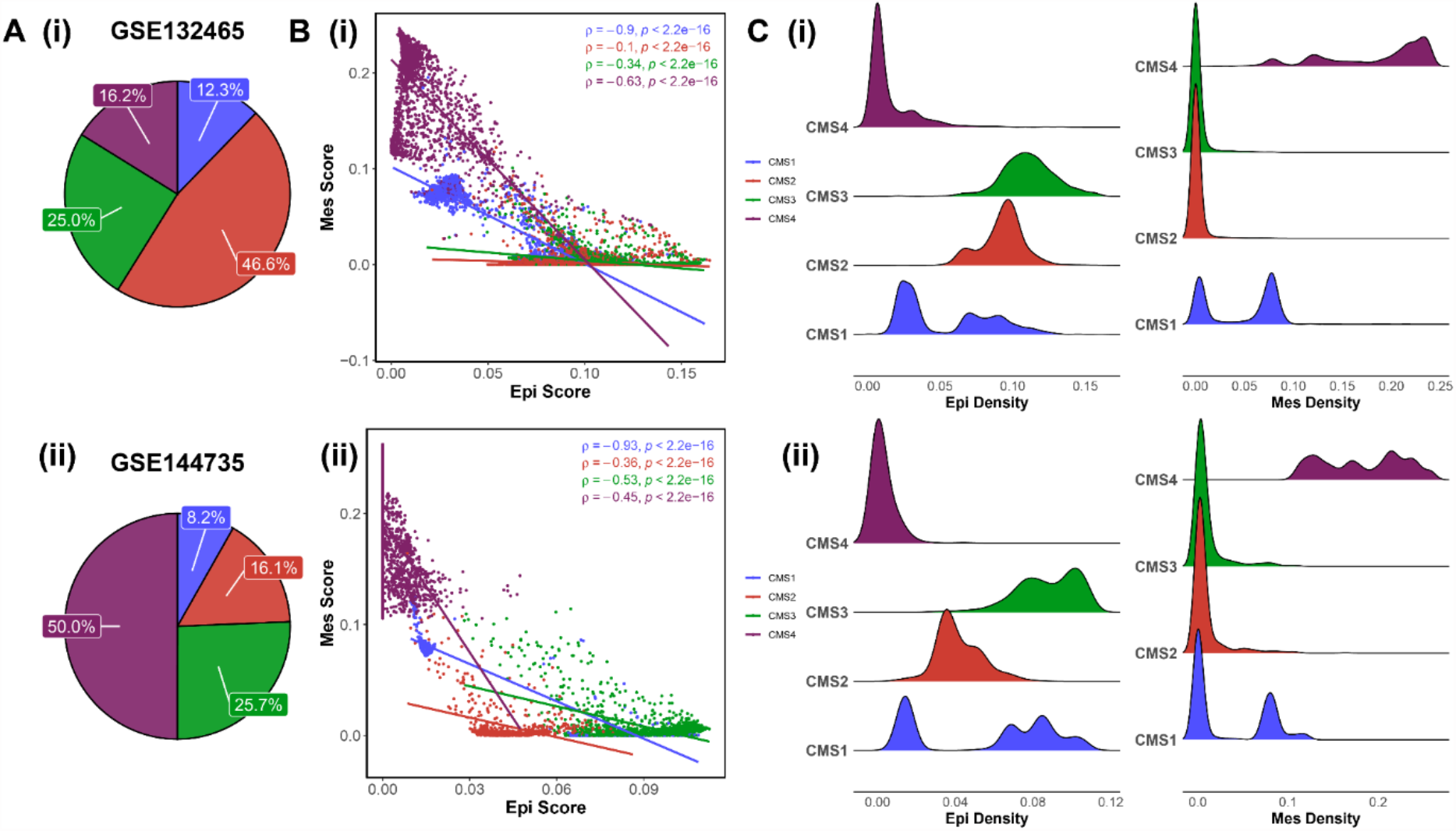
Heterogeneity among CMS subtypes along the epithelial-mesenchymal axis. **A) i)** Pie chart illustrating the percentage of cells classified as CMS1 (blue), CMS2 (red), CMS3 (green), and CMS4 (purple) in GSE132465. **B) i)** Scatter plot depicting the subtype-wise association between Epi (x-axis) and Mes scores (y-axis). Pearson’s correlation coefficient ‘ρ’ and p-values for subtype-wise correlation are shown. **C) i)** Subtype-wise kernel density estimate plots for ssGSEA scores of Epi (left) and Mes (right) signatures across cells in GSE132465. **A ii), B ii)**, and **C ii)** are the same as **A i), B i), C i)** respectively, but for GSE144735.

To assess how the epithelial and mesenchymal scores are distributed across the cells, we plotted the kernel density estimates of AUCell scores in a CMS subtype-wise manner. We noticed that the Epi density distribution for CMS4 shows a narrow peak centered around low Epi scores, whereas the CMS2 and CMS3 scores have wider distributions centered around higher Epi scores (**Fig 4C, i-ii, left)**. On the other hand, the CMS2 and CMS3 samples have low Mes scores with less variability (narrower peaks centered at low Mes scores), while CMS4 scores show higher average Mes scores as well as more heterogeneity in them (**Fig 4C, i-ii, right**). Interestingly, in both these datasets, CMS1 subtype distinctly showed two subpopulations in terms of their epithelial and mesenchymal scores (**Fig 4C, i-ii**). This bimodality was also seen in metabolic axes for the CMS1 subtype (**Fig S3, S4**). Together, our results support that CMS subtype comprises highly variable cells in terms of their E/M phenotype.

In both the scRNA-seq datasets, CMS specific associations of epithelial and mesenchymal programs with metabolic axes and PD-L1 were largely consistent with our earlier subtype-specific trend seen in bulk (TCGA) data. Simultaneously considering the associations between these axes of plasticity help in distinguishing similar CMS groups such as CMS1 and CMS4 (**Fig S3-S4**). For instance, In CMS1 cells, PDL1 and Epi scores are strongly negatively correlated, but not in other subtypes (**Fig S4D, i**).

Next, we wanted to quantify the amount of heterogeneity seen across these signatures in a subtype-specific manner. We used Shannon Entropy to calculate the variability among certain genes involved in a particular pathway across different sub-populations of CRC tumor cells (Conforte et al. 2019; Karolak et al. 2021). Higher entropy scores correspond to more variability in that axis for a particular cell. Cell-wise entropy values and ssGSEA scores largely showed a negative association with each other for all gene sets (**Fig 5, S5**). A possible explanation for this trend can be that once a cell acquires a particular phenotype, the genes involved in that pathway are coordinately being upregulated (or downregulated) and therefore have uniform high (or low) expression levels, reducing the underlying variability. Thus, it was unsurprising to notice the entropy for epithelial signature in CMS2 and CMS3 subtypes decreased with an increase in epithelial scores (**Fig 5A, i-ii**), potentially because those two subtypes are more epithelial relative to CMS1 and CMS4. Similarly, for CMS4, the most mesenchymal subtype, the increase in Mes scores associated with a decrease in entropy for mesenchymal signature in a cell (**Fig 5B, i-ii)**. Further, in CMS1, the subtype enriched in immune activation, an increase in PD-L1 signature scores correlated with a consistent decrease in entropy of corresponding signature (**Fig 5C, i-ii)**. However, the entropy of metabolic signatures did not show any CMS-specific trend, while they were also negatively correlated consistently with the corresponding signature scores (**Fig S5**).

**Figure 5:**
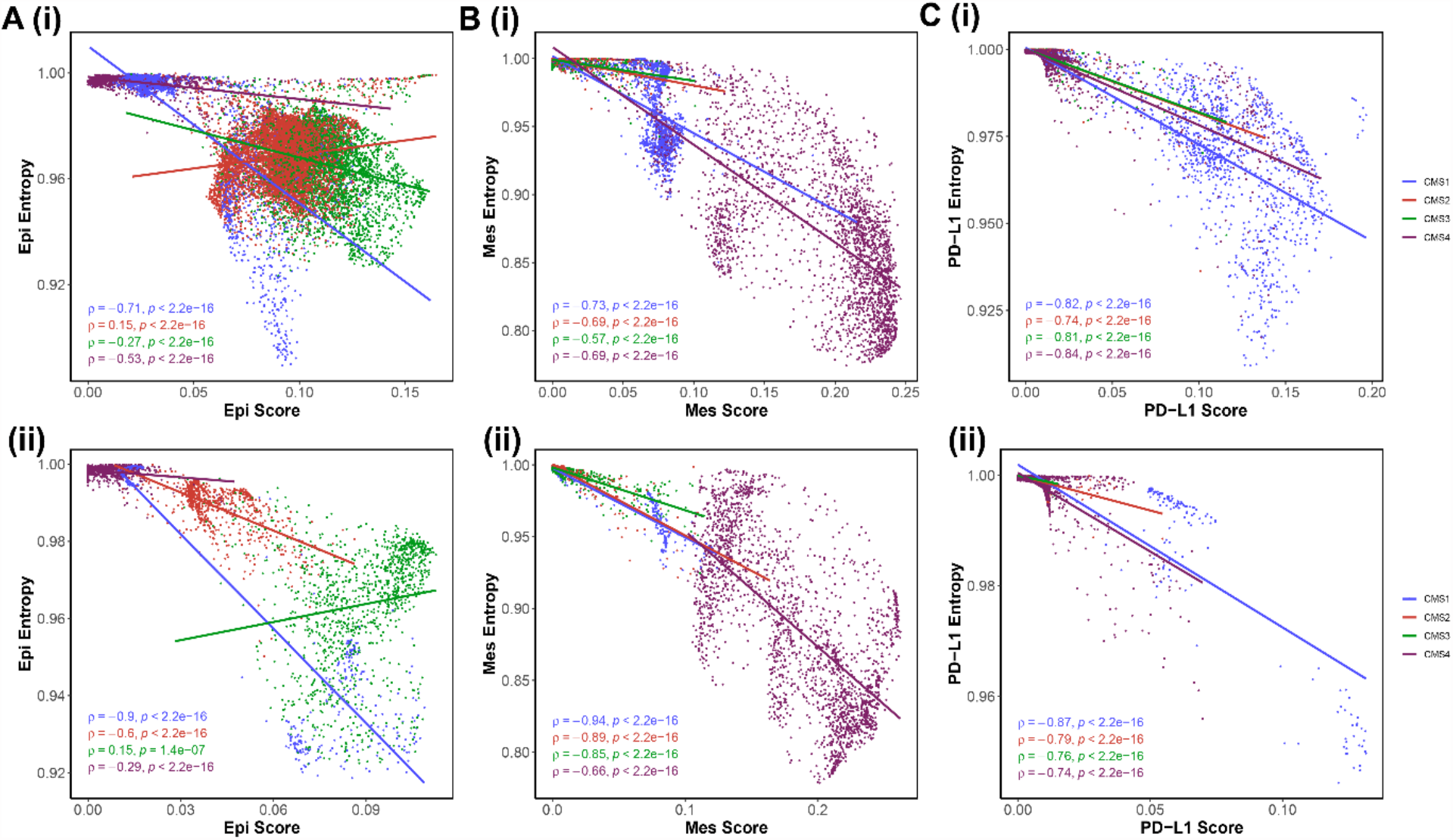
CMS-specific associations between cell-wise ssGSEA scores and entropy values of Epi and Mes genesets at a single-cell resolution. **A) i)** Scatter plot depicting the association between Epi scores (x-axis) and Epi entropy values (y-axis) for GSE132465. Same as **A i)** but for **B) i)** Mes score and entropy, **C) i)** PD-L1 score and entropy. **A ii), B ii)**, and **C ii)** are the same as **A i), B i)**, and **C i)** respectively, but for GSE144735. Pearson’s correlation coefficient ‘ρ’ and p-values for subtype-wise correlation are shown. Blue data points are for cells classified as CMS1, red for CMS2, green for CMS3, purple for CMS4.

## Discussion

Colorectal cancer is one of the most heterogeneous cancers characterized by intra and inter-tumoral heterogeneity. Thus, the classification of CRC into four CMS sub-types – while a helpful metric – does not entirely depict the heterogeneity in CRC. Here, we evaluated how some key axes that drive tumor progression and metastasis in carcinomas (immune evasion, metabolic reprogramming, and EMT) to understand their heterogeneity within CMS subtypes. Consistent with previous work in other cancer types, including ours, we observed a positive correlation between PD-L1 signature and mesenchymal state (Dongre et al. 2017; Muralidharan et al. 2022). However, across the CMS subtypes, PD-L1 signature score was not always negatively associated with epithelial scores, thus indicating that partial EMT state can also possess high immune-evasive traits (Dongre et al. 2021; Sahoo et al. 2021).

EMT is not binary switch but a spectrum of states including many epithelial/mesenchymal ones (Jolly et al. 2022; Subbalakshmi et al. 2022). This spectrum provides cancer cells stem-like traits, thereby facilitating aggressive tumor progression (Mani et al. 2008). Cell populations with higher plasticity along the E/M axis tend to be more metastatic and pose significant hurdles for treatment (Jolly et al. 2015, 2019). In this context, our analysis of CRC samples treated with EMT-inducers, SNAI1 and TGFβ revealed context-specific changes in the CMS state of samples with a concurrent alteration in their E/M state. While SNAI1, a highly specific EMT-TF reduces the epithelial and CMS3 enrichment and increases Mes and CMS4 scores, while TGFβ downregulates genes driving epithelial phenotype and increases enrichment of both CMS3 and CMS4 genes, possibly due to its role in mediating other axes of plasticity such as metabolic state (Shi et al. 2022). Our results showing CMS4 to be most mesenchymal are consistent with earlier observations about higher methylation of miR-200 family in CMS4 cell lines and tumors (Fessler et al. 2016). Further, transcriptomes of TGFβ treatment of CRC organoids resemble the CMS4 signature seen in human tumors (Flum et al. 2022).

Our results for scRNA-seq and bulk RNA-seq analysis are largely self-consistent across the CMS subtypes, such as antagonism between epithelial and mesenchymal programs, or the association of those programs with metabolic axes and PD-L1 signature enrichment scores. The different modalities of associations observed between these axes may explain the CMS subtype-specific observations of patient survival and/or sensitivity to various therapeutics and remains a key focus for our future work.

A key point that our scRNA-seq analysis reveals is that CMS2 and CMS3 are relatively most epithelial, while CMS4 and CMS1 being more mesenchymal. This categorization is reminiscent of previous observations such as the multi-omics profiling of CRC cell lines suggesting that CMS2 and CMS3 ones are more colon-like, while CMS and CMS4 ones are more undifferentiated and had higher expression of genes associated with EMT and TGFβ signaling (Berg et al. 2017). Similarly, in TCGA CRC data, most patients from ZEB1^hi^ group belonged to CMS4 subtype, while the ZEB1^lo^ group was mainly composed of CMS2 and CMS3 tumors (Xu et al. 2022). CMS4 subtype expression also correlates well with the signature of EpCAM^lo^ sub-population in HCT116 and SW480 cells (Sacchetti et al. 2021).

Besides EMT, metabolic reprogramming is another key axis of cancer cell plasticity. Recent studies have shown that along with classical Warburg effect observed in cancer cells, some cancers, including cervical and breast cancer, predominantly use OXPHOS as a primary energy source (Rodríguez-Enríquez et al. 2010; Hernández-Reséndiz et al. 2015). CRC cells have been reported to have a higher OXPHOS rate compared to normal colon cells (Kaldma et al. 2014). Through a mechanism known as Reverse Warburg effect, elements of tumor microenvironment, such as cancer-associated fibroblasts (CAF), can regulate the OXPHOS-glycolysis metabolic switch in cancers (Bonuccelli et al. 2010). Thus, varying microenvironments may possibly explain our counterintuitive association of glycolysis with epithelial and mesenchymal scores. Metabolism-based characterization of CRC samples has also been attempted recently (Zhang et al. 2020). Future efforts enabling more accurate classification of CRC patients into different subgroups with specified vulnerability can integrate such efforts being made to unravel phenotypic heterogeneity among multiple interconnected axes of plasticity.

## Author contributions

MKJ designed and supervised research and obtained funding. MS, SR, JMV, YRG, SM and SV performed research. MS and SR analyzed data and wrote the first manuscript draft. All authors have read and agreed to submission of this version of the manuscript.

## Funding

This work was supported by Ramanujan Fellowship (SB/S2/RJN-049/2018) awarded to MKJ by Science and Engineering Research Board (SERB), Department of Science and Technology, Government of India.

## Conflict of Interest

The authors declare no conflict of interest.

## Supplementary Figures

**Figure S1:**
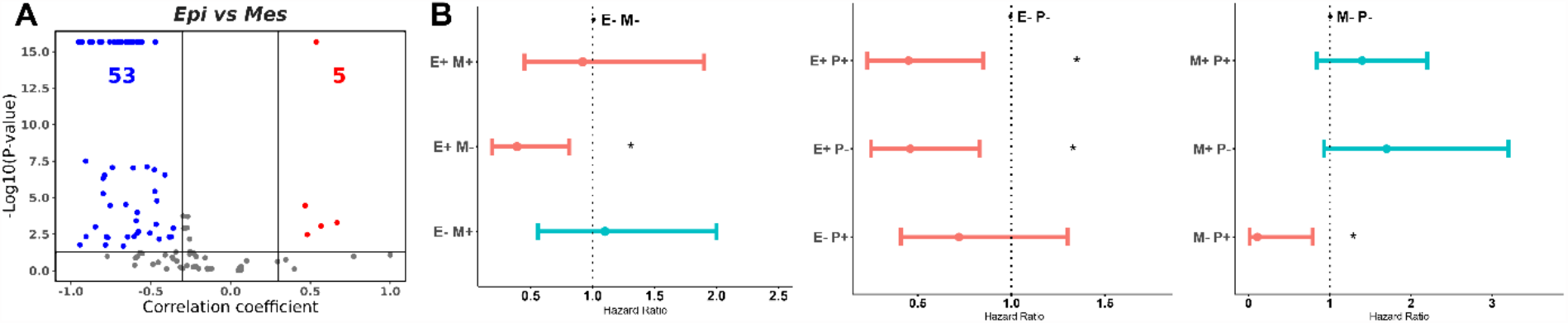
**A)** Volcano plot depicting Spearman correlation coefficient (x-axis) and -log10(p-values) (y-axis) for Epi vs. Mes scores. Boundaries for significant correlation are set at R > ± 0.3 and p < 0.05. Red data points indicate datasets for which the association is significantly positive, blue for negative, and gray for insignificant correlation. **B)** Forest plots depicting mean hazard ratios (HR) ± 95% confidence intervals and corresponding p-values (‘*’ for p <0.05) for overall survival associated with concurrent enrichment of epithelial and mesenchymal (left), epithelial and PD-L1 (middle) and mesenchymal and PD-L1 signatures (right). Mean HR values > 1 are shown in blue while those < 1 are shown in red. (+) and (-) subgroups are based on median values.

**Figure S2:**
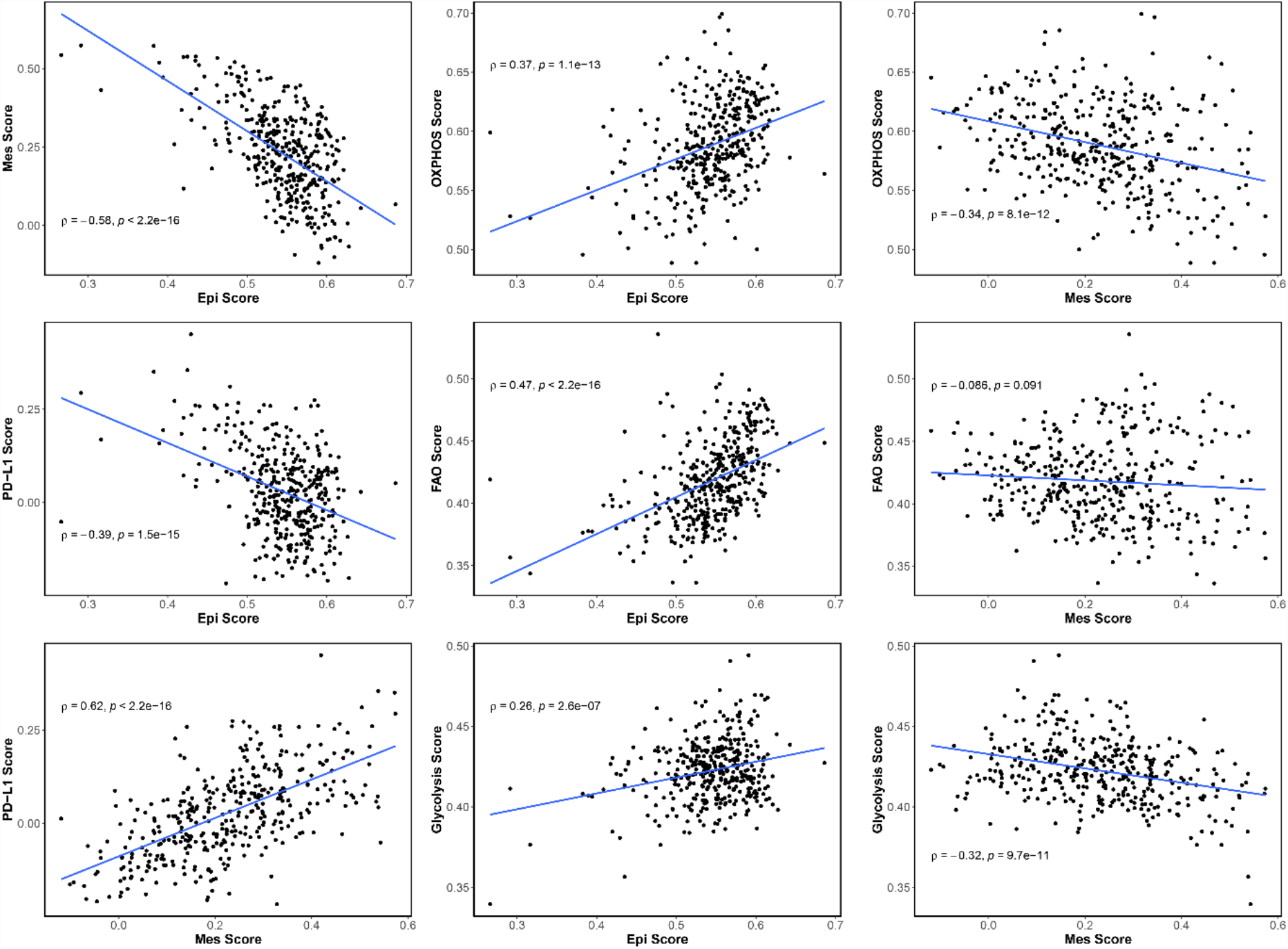
Scatter plots illustrating associations between Epi vs. OXPHOS, FAO, glycolysis and PD-L1 scores and for Mes vs. OXPHOS, FAO, glycolysis and PD-L1 scores for TCGA colorectal cancer patient samples.

**Figure S3:**
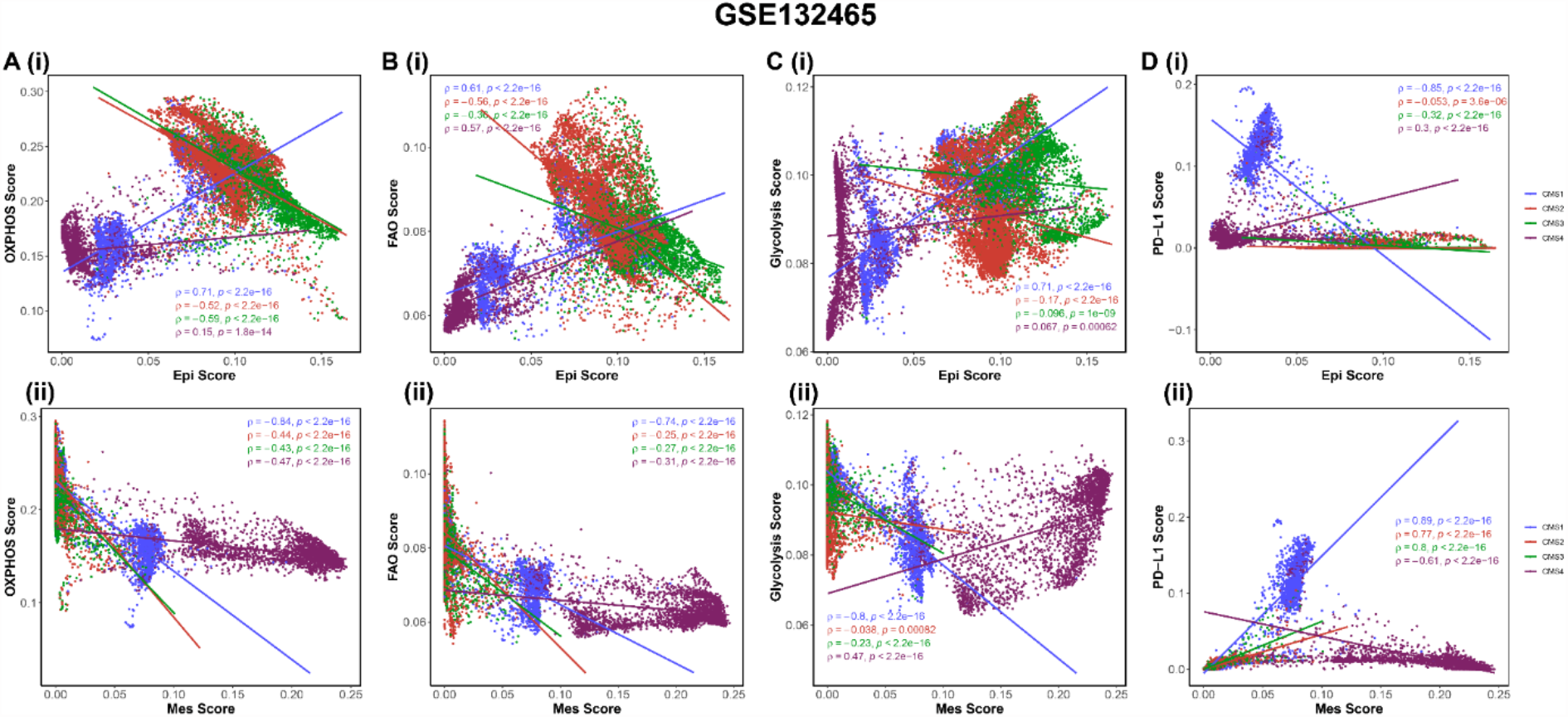
CMS-specific associations of epithelial and mesenchymal programs with metabolic axes and PD-L1 in GSE132465. **A (i)** Scatter plot depicting association between Epi (x-axis) and OXPHOS scores (y-axis). **A (ii)** Same as **A (i)** but for Mes and OXPHOS scores, **B (i)** Epi and FAO, **B (ii)** Mes and FAO, **C (i)** Epi and Glycolysis, **C (ii)** Mes and Glycolysis, **D (i)** Epi and PD-L1, and **D (ii)** Mes and PD-L1 scores. Pearson’s correlation coefficient ‘ρ’ and p-values for subtype-wise correlation are shown. Blue data points are for cells classified as CMS1, red for CMS2, green for CMS3, purple for CMS4.

**Figure S4:**
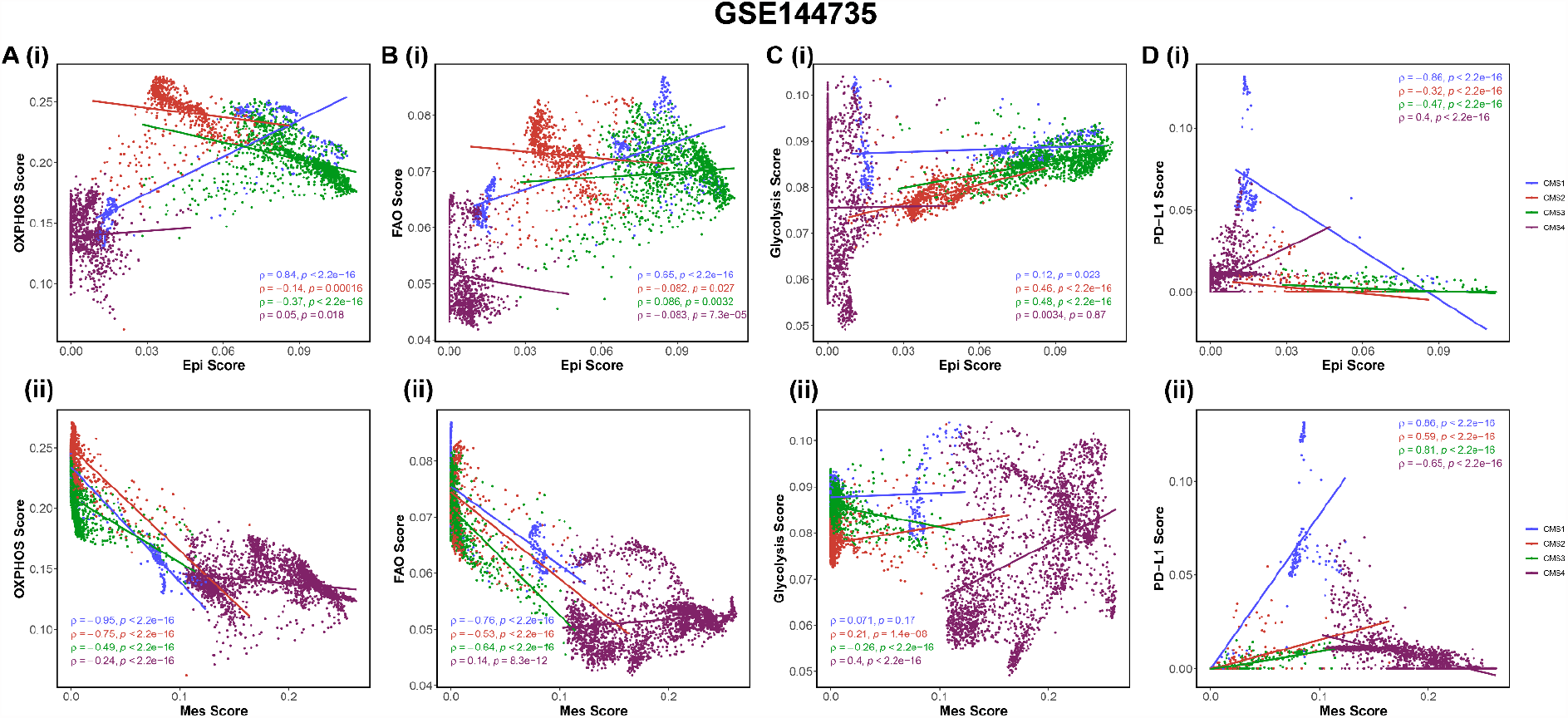
CMS-specific associations of epithelial and mesenchymal programs with metabolic axes and PD-L1 in GSE144735. **A (i)** Scatter plot depicting association between Epi (x-axis) and OXPHOS scores (y-axis). **A (ii)** Same as **A (i)** but for Mes and OXPHOS scores, **B (i)** Epi and FAO, **B (ii)** Mes and FAO, **C (i)** Epi and Glycolysis, **C (ii)** Mes and Glycolysis, **D (i)** Epi and PD-L1, and **D (ii)** Mes and PD-L1 scores. Pearson’s correlation coefficient ‘ρ’ and p-values for subtype-wise correlation are shown. Blue data points are for cells classified as CMS1, red for CMS2, green for CMS3, purple for CMS4.

**Figure S5:**
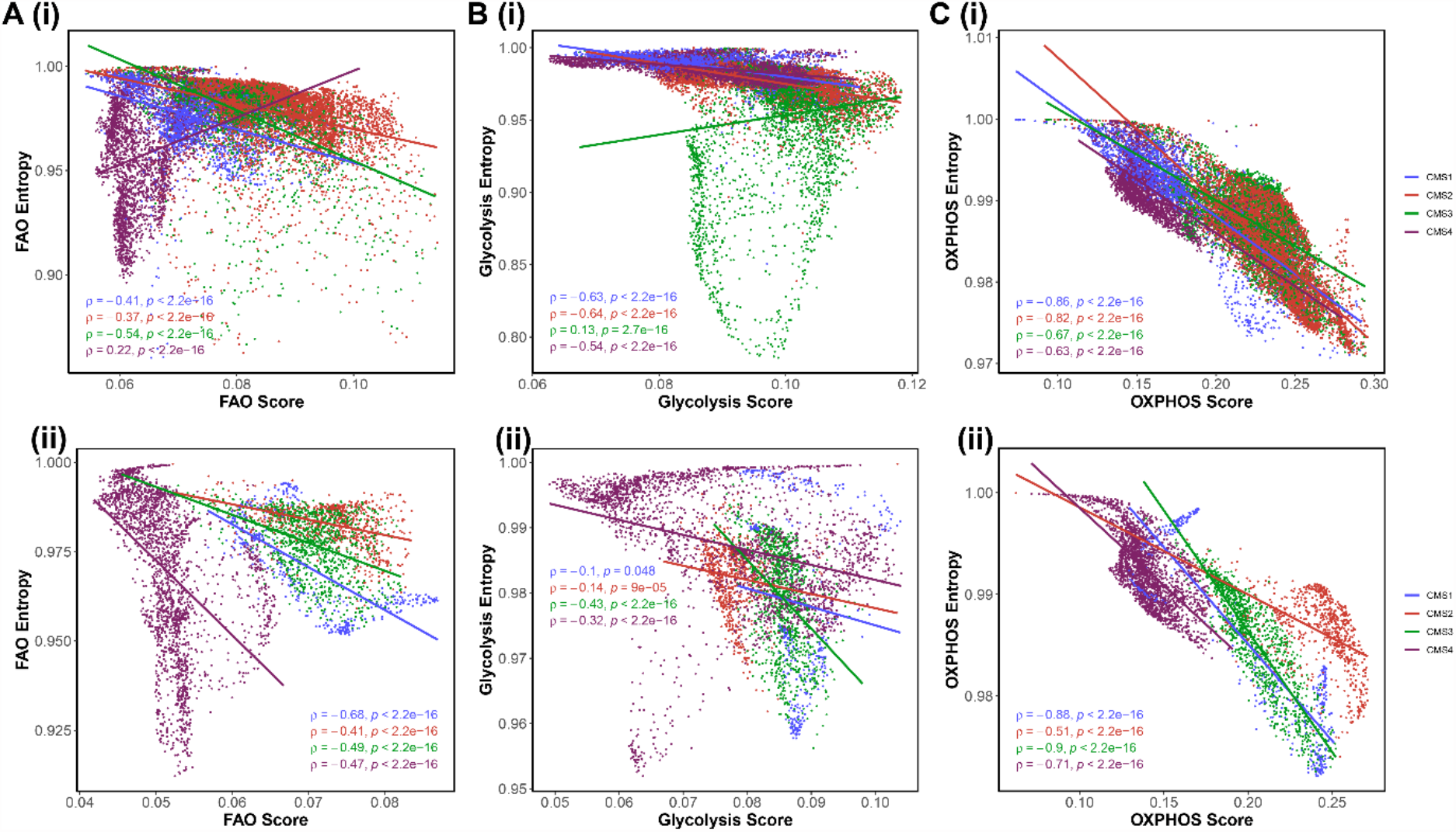
CMS-specific associations between cell-wise ssGSEA scores and entropy values of metabolic and PD-L1 gene sets at a single cell resolution. **A (i)** Scatter plot depicting association between FAO scores (x-axis) and FAO entropy values (y-axis) for GSE132465. Same as **A (i)** but for **B (i)** Glycolysis and **C (i)** OXPHOS. **A (ii), B (ii)** and **C (ii)** are the same as **A (i), B (i)** and **C (i)** respectively, but for GSE144735. Pearson’s correlation coefficient ‘ρ’ and p-values for subtype-wise correlation are shown. Blue data points are for cells classified as CMS1, red for CMS2, green for CMS3, purple for CMS4.

## Supplementary Tables

**Table S1**

Description of 101 bulk transcriptomic datasets used in this study along with Spearman’s Correlation coefficient ‘R’ and corresponding p-values for correlation of Epithelial and Mesenchymal signature with PD-L1 gene signature and metabolic signatures (glycolysis, FAO and OXPHOS).

**Table S2**

Gene sets used for scoring the CMS subtypes, EMT, metabolism and PD-L1 gene signatures.

## Notes

### Competing Interest Statement

The authors have declared no competing interest.

